# Immune DNA signature of T-cell infiltration in breast tumor exomes

**DOI:** 10.1101/047753

**Authors:** Eric Levy, Rachel Marty, Valentina Garate-Calderon, Brian Woo, Michelle Dow, Ricardo Armisen, Hannah Carter, Olivier Harismendy

**Affiliations:** Division of Biomedical Informatics, Department of Medicine, University of California San Diego; Bioinformatics and Systems Biology Graduate Program, University of California San Diego; Division of Medical Genetics, Department of Medicine, University of California San Diego; Centro de Investigación y Tratamiento del Cancer, Facultad de Medicina, Universidad de Chile; Center for Excellence in Precision Medicine, Pfizer Chile; Moores Cancer Center, University of California San Diego; Institute for Genomic Medicine, University of California San Diego

**Author notes:** Correspondence to: (*) Olivier Harismendy –.

## Abstract

Tumor infiltrating lymphocytes (TILs) have been associated with favorable prognosis in multiple tumor types. The Cancer Genome Atlas (TCGA) represents the largest collection of cancer molecular data, but lacks detailed information about the immune environment. Here, we show that exome reads mapping to the complementarity-determining-region 3 (CDR3) of mature T-cell receptor beta (*TCRB*) can be used as an immune DNA (iDNA) signature. Specifically, we propose a method to identify CDR3 reads in a breast tumor exome and validate it using deep *TCRB* sequencing. In 1,078 TCGA breast cancer exomes, the fraction of CDR3 reads was associated with TILs fraction, tumor purity, adaptive immunity gene expression signatures and improved survival in Her2+ patients. Only 2/839 *TCRB* clonotypes were shared between patients and none associated with a specific HLA allele or somatic driver mutations. The iDNA biomarker enriches the comprehensive dataset collected through TCGA, revealing associations with other molecular features and clinical outcomes.

---

In breast cancer, the presence of tumor infiltrating lymphocytes (TILs), and more specifically T-lymphocytes, is associated with good survival^1,2^ and response to neo-adjuvant treatment^3,4^. The different breast cancer subtypes do not significantly differ in fraction of TILs, which is relatively low^5^, but this metric has prognostic or predictive value in triple negative breast cancer (TNBC) and Her2+ breast cancer^4,6,7^. In order to further distinguish the different cell type populations, other studies have used immunohistochemistry to detect cell surface markers (e.g. CD3, CD8, CD20), demonstrating, for example, that the predictive value of B-cell infiltration is independent of cancer subtype or other clinical factors^8^, or that CD8+ T-cell infiltration is of good prognosis in basal TNBC^5^. A related clinical-grade assay, the immunoscore, is being proposed for colorectal cancer^9^, but requires further evaluation in breast cancer^3^.

Analysis of gene expression signatures can also be used to infer the presence of immune cells and their role in immune signaling within the tumor microenvironment. High levels of a TIL-associated signature is associated with good prognosis in ER-breast cancer^10^. Gene expression signatures specific to T-cells^5,11^ and B-cells^12^ also have prognostic or predictive value in specific cancer subtypes. Interestingly, while the expression of metagenes is not different between breast cancer subtypes, their prognostic significance varies. For example, the expression of a T-cell metagene is associated with good prognosis in ER-or Her2+ tumors^11^. More recently, the gene expression measurements in heterogeneous tumor samples have been deconvolved using machine learning to determine the relative abundance of up to 22 immune cell types^13^. This association revealed an opposite survival association of plasma cells and neutrophils^14^.

Correlations have been observed between the extent of T-cell infiltration and clinical prognosis in breast cancer subtypes. However, this effect is indirect, related to the T-cells’ role in tumor control and is dependent on their tumor reactivity. Thus a deeper characterization of the T-cell repertoire can provide more information about its diversity, the associated tumor reactivity, and antigen specificity. Recent technical progress has enabled the characterization of T-cell repertoires by deep sequencing of the VDJ rearrangement at the complementarity determining region 3 (CDR3) of *TCRB*^15^, and has been used to observe at an unprecedented resolution the clonal diversity of T-cells during infection and in solid tumors^16–18^. Deep repertoire sequencing performed in tumors of the colon^18^, ovary^19^, kidney^20^, pancreas^21^, or lung^22^ have addressed methodological challenges and have confirmed the diversity and specific landscape of TILs. However, the technical validity and clinical utility of TCR repertoire characterization in tumors remains to be established. In particular, it is not yet clear whether the quantity (fraction of T-cells) or the diversity (relative abundance of specific clones) is more important to predict disease progression and response to treatment. Similarly, we do not know the extent of clonotype sharing between patients or between tumor, lymph nodes, and metastasis of the same patient or whether any clinical association with these patterns can be determined. Overall, the understanding of the tumor immune environment remains fragmented, and a more comprehensive integrated approach is needed to characterize the tumor immune landscape, as recently suggested by the colorectal cancer anti-genome study^23^. Comprehensive profiling of the immune environment, including T-cell repertoire, needs to be expanded to larger, well-annotated cohorts to establish its potential utility. The Cancer Genome Atlas (TCGA) provides a large resource of molecular data that can be interrogated for this immune environment^24^. Here, we show that it is possible to re-analyze tumor exomes and transcriptomes from TCGA to quantify and characterize infiltrating T-cells through the detection of a rearranged CDR3 of the *TCRB* gene. We first establish the feasibility of the approach by characterizing the rearranged TCR repertoire using deep sequencing of a breast cancer specimen and comparing the resulting clonotypes to the ones identified in the whole exome sequence of the same sample. We then identify CDR3 reads in TCGA breast cancer tumors, and show their correlation with other markers of immune infiltration. We further evaluate their prognostic value in breast cancer subtype and investigate clonotype diversity and sharing between patients and specimens.

## Results

### Deep TCR repertoire sequencing

We sequenced the repertoire of three triple negative breast cancer (TNBC) samples selected for their variable TIL· contents. Two samples had a high amount of infiltration (45% and 40%), and one sample was chosen as a negative control (0%). Starting from 5 μg of DNA (~8x10^5^ total cells), we identified between 15×10^3^ and 30×10^3^ CDR3 rearrangements per tumor (Supplementary Fig. 1A). Interestingly, even the tumor sample with no histological evidence of TILs shows multiple rearrangements, suggesting a limitation of histological evaluation using a selected tissue section. Thanks to internal standards, the assay was able to precisely estimate the abundance of each clone and the overall clonality of each sample. The most clonal sample (OX1285: clonality=0.22) contained the most abundant clone at 8% prevalence. In contrast, the two other samples had clonalities of 0.15 and 0.09, and the most abundant clone at 1.7% each. The abundance of each clone was highly reproducible between two adjacent tissue sections (r=0.99), suggesting a local homogeneity of the T-cell population (Fig. 1A). In complement to this data generation, we also evaluated the feasibility of using archival FFPE specimens for deep *TCRB* amplicon sequencing. Two samples showing the most fragmented DNA (average size <1.1 kb) had poor overall *TCRB* representation when compared to the matched frozen. The least fragmented sample had the most reproducible results when compared to a matched frozen with an overall underestimate of the absolute clonotype frequency (Supplementary Fig. 1B-D) This demonstrates that by using stringent DNA sample quality control, archival samples may be used for deep repertoire sequencing, albeit resulting in reduced accuracy.

**Figure 1:**
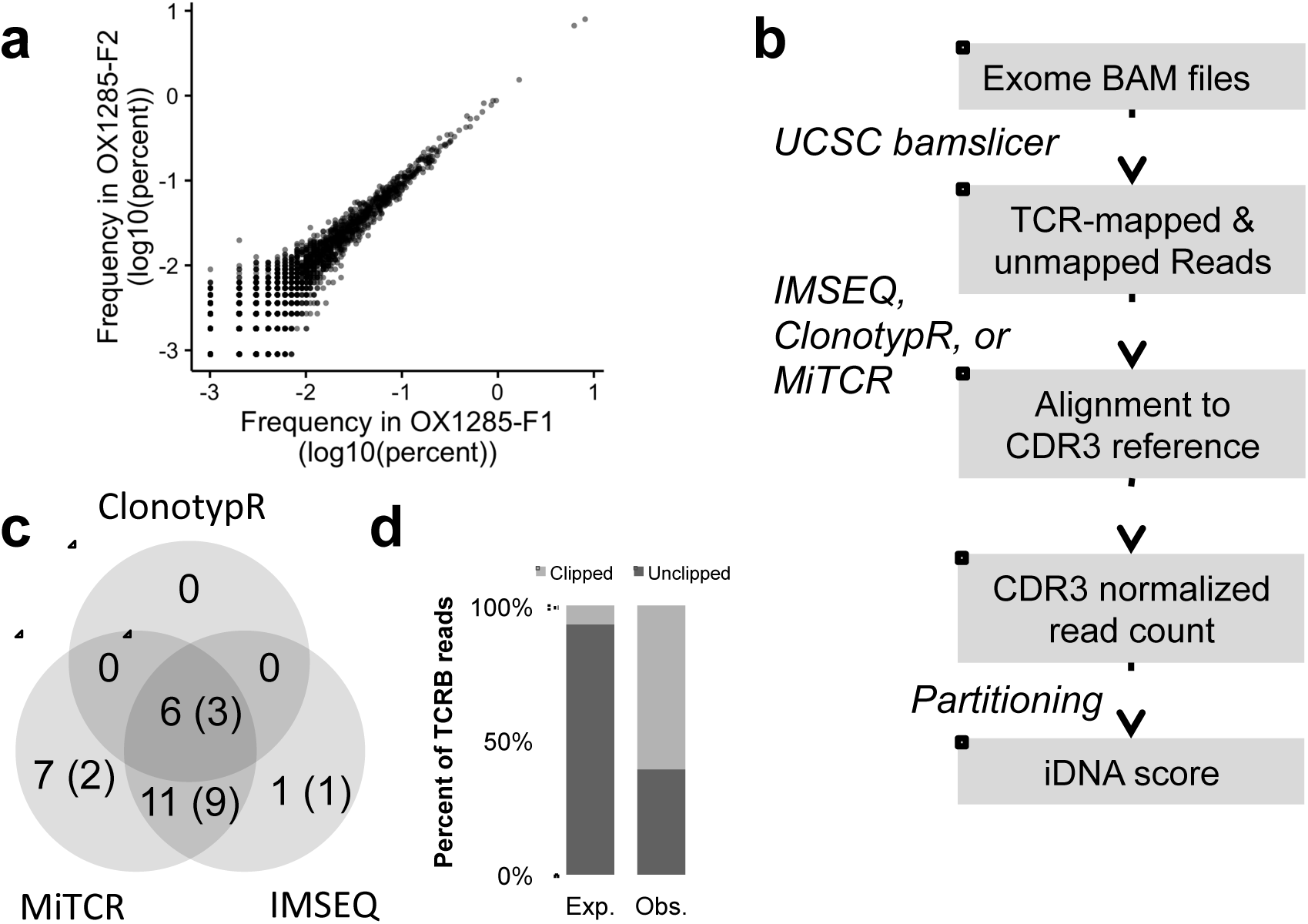
Identification of CRD3 reads in whole-exome data. **(a)** Clonotype abundance determined by deep repertoire sequencing (ImmunoSeq) in two adjacent breast cancer tissue sections, **(b)** Workflow to extract and identify rearranged CDR3 reads from exome datasets, **(c)** Comparison of the number of CDR3 reads identified by each clonotyping tools. The number in parenthesis indicates the subset of clonotypes also identified by deep repertoire sequencing, **(d)** Fraction of clipped reads mapped to the TCR region in the exome BAM file. The expected is estimated from all mapped reads in the exome.

Identification of CDR3 reads in tumor exomes Sequencing a full, deep repertoire of TILs is costly and requires large amounts of DNA to ensure that sufficient clonal diversity is being captured. We thus sought to determine whether any of the *TCRB* clonotypes could be identified in exome sequencing data, which would permit the use of public cancer genomic data. Indeed, most exome capture kits contain probes overlapping the V and J genes of the *TCRB* locus. While such probes have been designed to capture the naïve TCR region, it is likely that a rearranged DNA fragment can be captured if it has sufficient overlap with the reference sequence to allow probe hybridization. To test this hypothesis, we sequenced 205×10^6^ reads from the exome of sample OX1285, for which we obtained deep repertoire data (Supplementary Table 1). Of these, 784×10^3^ reads did not map to the reference genome and 241×10^3^ mapped to the reference *TCRB* locus. In order to identify reads mapping to a rearranged CDR3 domain of *TCRB* (referred to as CDR3 reads), we benchmarked three different tools: clonotypR^25^, IM-SEQ^26^ and MiTCR^27^ (Fig. 1B), each originally designed to analyze deep repertoire sequencing experiments. Each tool identified between 10 and 38 reads assigned to a CDR3 (Table 1). Across all three methods, we identified a total of 26 clonotypes, 15 of which were present in the deep repertoire dataset (Fig. 1C). Interestingly, 60% of the CDR3 reads mapped imperfectly (clipped reads) to the reference *TCRB* locus (Fig. 1D), consistent with their mature *TCRB* origin and suboptimal alignment to the naïve *TCRB* genes Fourteen clonotypes were identified by two or more methods. ClonotypR was the most stringent, only finding 6 clonotypes, all identified by the other tools. In contrast, MiTCR was the most lenient, with 7 unique clonotypes, 2 of which were present in the deep repertoire. Overall, IMSEQ offered the best compromise between sensitivity – 72% present in deep sequencing – and specificity – 94% shared with another tool – and was used for the rest of the analysis. The fraction of CDR3 reads detected by IMSEQ is 0.09 reads per million reads (RPM) sequenced. Interestingly, assuming that this tumor had 20-40% of infiltrating T-cells, this value was consistent with the order of magnitude estimated by simulations (~10^‐1^ – Supplementary Fig. 2 and Methods). The same simulation also suggested that, at typical exome sequencing coverage depth (100 fold), CDR3 reads could be detected in tumors with more than 3% T-cell infiltration. These results provide evidence that genuine CDR3 reads can be identified in exome sequencing data from a bulk tumor.

**Table 1:** Distribution of clonotypes identified in OX1285 exome using three CDR3 detection tools. (*) indicates rescued out-of-frame CDR3 reads in IMSEQ.

| Clone ID | Number of reads supporting each clone in the exome data |  |  | ImmunoSeq Abundance (%) |
| --- | --- | --- | --- | --- |
|  | ClonotypeR | IMSEQ | MiTCR |  |
| c1 | 1 | >0* | 6 | 8.10 |
| c2 | 0 | 0 | 1 | 1.67 |
| c3 | 0 | 1 | 1 | 0.89 |
| c4 | 0 | 2 | 2 | 0.71 |
| c5 | 0 | 1 | 1 | 0.52 |
| c6 | 0 | 2 | 2 | 0.24 |
| c7 | 0 | 1 | 1 | 0.21 |
| c8 | 0 | 1 | 1 | 0.13 |
| c9 | 0 | 1 | 0 | 0.12 |
| c10 | 0 | 1 | 1 | 0.06 |
| c11 | 0 | >0* | 1 | 0.04 |
| c12 | 1 | 1 | 1 | 0.03 |
| c13 | 3 | >0* | 3 | 0.02 |
| c14 | 0 | >0* | 1 | 0.01 |
| c15 | 0 | 0 | 1 | 0.003 |
| c16 | 2 | 2 | 0 | NA |
| c17 | 0 | 1 | 1 | NA |
| c18 | 0 | 0 | 1 | NA |
| c19 | 0 | 0 | 1 | NA |
| c20 | 0 | 0 | 1 | NA |
| c21 | 0 | 0 | 1 | NA |
| c22 | 1 | 1 | 1 | NA |
| c23 | 0 | >0* | 2 | NA |
| c24 | 0 | 0 | 2 | NA |
| c25 | 2 | 2 | 2 | NA |
| c26 | 0 | 0 | 3 | NA |

### Identification of CDR3 reads in the TCGA breast cancer exomes

Using the approach validated above, we analyzed the exome sequences of 1078 breast cancer tumors characterized through TCGA. We identified CDR3 reads in 473/1,078 (44%) tumors (Supplementary Table 2). For some of the downstream analysis, we smoothed the normalized CDR3 read content of each tumor into an immune DNA (iDNA) score: 0 for absence of CDR3 reads, and 1-10 for the increasing deciles of the distribution of normalized CDR3 read count (CDR3 RPM). CDR3 RPM was associated with high TILs (p<3×10^-7^ – Wilcoxon test). Indeed, only 19% of the tumors with no CDR3 reads (iDNA=0) had more than 5% TILs, in contrast to 49% of the tumors with an iDNA score of 10 (Fig. 2A). Importantly, TIL measurements refers to total TILs, not only T-cells, and this measurement may vary between sample collection sites and pathologists, despite efforts to standardize it^3^. For a more quantitative evaluation, we chose to compare the fraction of tumor CDR3 reads to the tumor molecular purity^28^. We observed that the fraction of CDR3 reads was inversely correlated (r=-0.39) with tumor purity, with 43% of tumors without CDR3 reads having purity higher than 80%, in contrast to only 4% of the tumors with an iDNA score of 10 (Fig. 2B). These results suggest that the CDR3 reads identified in the tumor exome truly originate from T-cells, and that their relative abundance is directly associated with the fraction of infiltrating lymphocytes.

**Figure 2:**
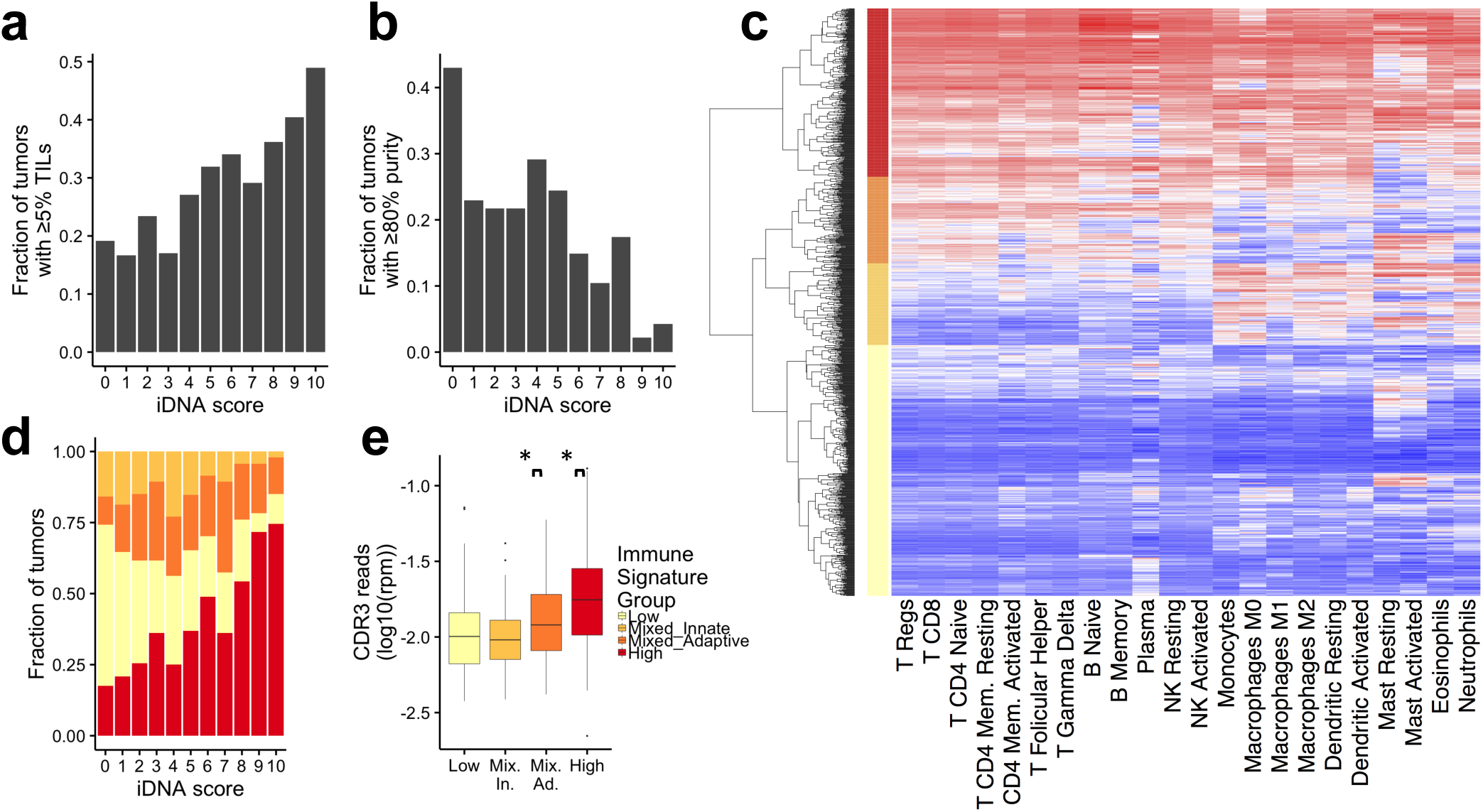
Association between iDNA score and the tumor immuneenvironment. **(a)** Fraction of tumors with more tha? 5% TILs in each iDNA score. **(b)** Fraction of tumors with more tha? 80% tumor purity in each iDNA score, **(c)** Clustering of 1072 breast tumors according to the ssGSEA score (red:high, blue:low) of 22 immune gene signatures. The four main clusters, high (red), mixed adaptive (dark orange), mixed innate (light orange) and low (yellow) are labeled on the y-axis. **(d)** Distribution of tumors between the four immune signature groups with increasing iDNA scores, **(e)** Distribution of the CDR3 reads normalized abundance in tumors of the four immune signature groups (*) p<0.01, t-test. Only tumors with CDR3 reads are included.

### iDNA score correlates with adaptive immunity expression signatures

In order to further explore the variation in iDNA scores, we used the level of gene expression to measure the relative enrichment for 22 immune cells signatures in each tumor^13^. Expression signatures of different immune cells are often correlated, but using unsupervised clustering of the signature scores, we were able to distinguish 4 groups of tumors: immune low (n=458), mixed-adpative (n=159), mixed-innate (n=149) and high (n=307) (Fig. 2C). Seventy five percent of the tumors in the immune-low group did not have CDR3 reads (iDNA score=0, Fig. 2D). In contrast, 65% of the immune-high tumors had CDR3 reads (iDNA score>0). The immune-high group showed high levels of both adaptive and innate signatures. In contrast, the immune-mixed groups showed a clear distinction in activity levels of adaptive and innate immune cells. For tumor with iDNA scores greater than 0, the CDR3 RPM was higher in mixed-adaptive than mixed-innate groups, and the latter was not different from the immune-low group (Fig. 2E). These results indicate that the abundance of CDR3 reads in exomes is correlated with known expression signatures of adaptive immunity.

### CDR3 read content associates with survival

The fraction of tumors positive for iDNA was not different between breast cancer subtypes (Fig. 3A) We show, however, that a positive iDNA score was associated with better overall survival (HR=3.17 [1.18-8.51], p=0.022) in Her2+ breast cancer, but not in hormone positive, Her2 negative (HR+/Her2-) or TNBC (Fig. 3B and Supplementary Fig. 3). Most Her2+ patients in the TCGA cohort were likely treated with anti-Her2 antibody therapy. Therefore, the iDNA score is predictive rather than prognostic for the Her2+ subtype. Interestingly, the fraction of TILs alone was not predictive of response in Her2+ tumors (Fig. 3C), suggesting the superiority of a DNA based measurement of mature T-cell content over the histological estimate of lymphocyte content.

**Figure 3:**
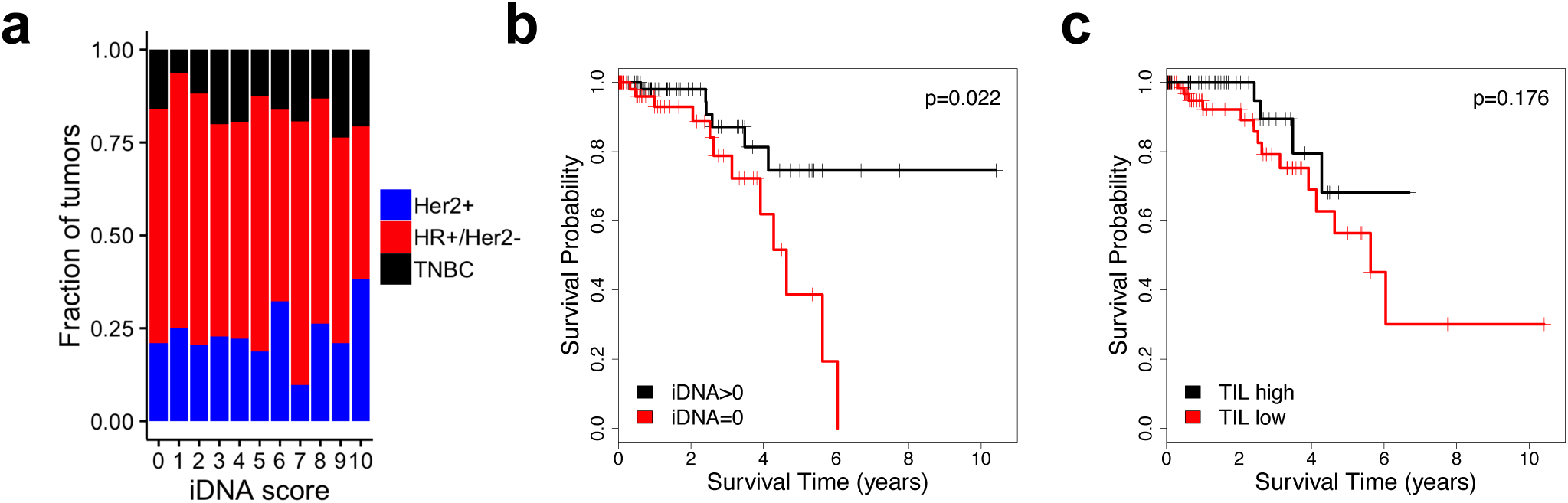
Association with survival. **(a)** Distribution of tumors by breast cancer histological subtypes at increasing iDNA scores, **(b-c),** Kaplan-Meier survival analysis of Her2+ patients as a function of iDNA score (b) and TIL content (c) with significance of the Cox proportional hazard ratio. Hazard ratio is 3.17 [1.18-8.51], p=0.022 for iDNA, and 0.462 [0.151-1.142], p=0.176 for TIL.

### *TCRB* expression and clonal diversity and sharing

We then asked whether the CDR3 sequences identified in the tumor exome were expressed and how they relate to the overall expression of the *TCRB* gene. Of the 1,074 tumor specimens with available transcriptome data, we were able to identify CDR3 reads in 906 (84%) of them. The fraction of CDR3 reads in the RNA is correlated with the overall *TCRB* expression level (including non-CDR3 reads - r=0.40 p<10^-16^ – Fig. 4A). There were 435 tumors with evidence of CDR3 reads in both tumor DNA and RNA, and the fraction of CDR3 reads in the RNA and DNA was correlated (r=0.33, p= 6.304×10^-13^, Fig. 4B). Interestingly, the overall expression of the *TCRB* gene increased from tumors with no CDR3 in RNA nor in DNA (N=132), CDR3 reads in DNA only (N=36), in RNA only (N=471) or in both (N=435 Fig. 4C p<0.001 - ANOVA). This observation suggests that some tumors may have few infiltrating T-cells (exome CDR3 negative), but these T-cells express sufficient levels of *TCRB* for the CDR3 to be detected in the transcriptome. Conversely, a few tumors display unambiguous T-cell infiltration (exome CDR3 positive), but expression of the *TCRB* gene is too low to detect CDR3 sequences in the transcriptome. This result underscores the importance of studying T-cell infiltration by DNA or histology based methods for a more quantitative assessment of their level of infiltration, in contrast to RNA-based methods, which can be confounded by the regulation of the *TCRB* gene expression.

**Figure 4:**
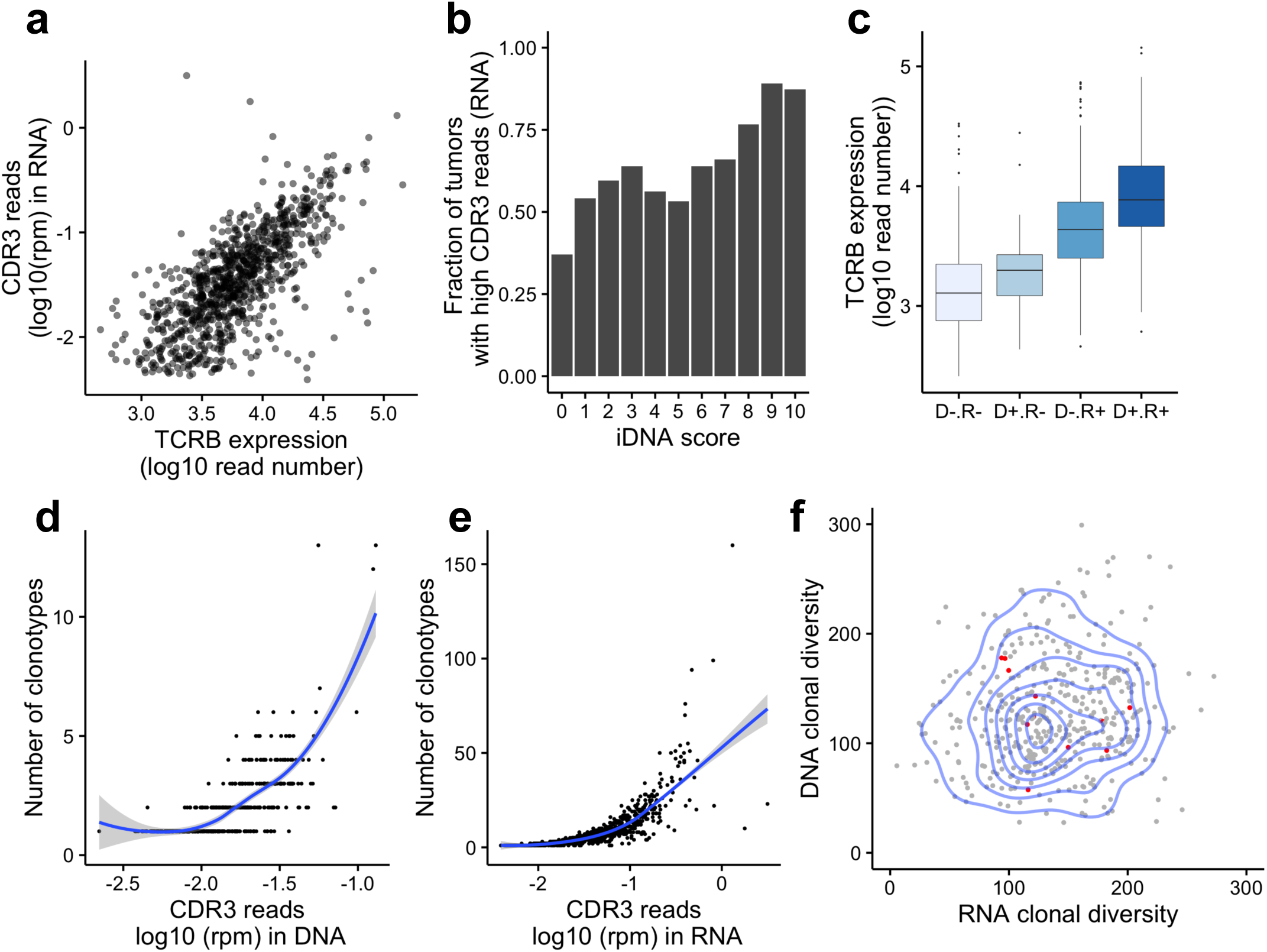
TCRB expression and clonal diversity. **(a)** Correlation between overall TCRB expression (x-axis) and CDR3 read abundance in RNA (y-axis), r=0.69. **(b)** Distribution of the normalized CDR3 read count from RNA-seq at each iDNA score. **(c)** Distribution of the TCRB expression level in groups of tumors where CDR3 reads can be identified in neither DNA nor RNA (D-R-), DNA only (D +R-), RNA only (D-R+) or both (D+R+) (p<0.001 by ANOVA). **(d)** The number of clonotypes identified increases with the fraction of CDR3 reads in the exome dataset (x-axis). **(e)** The number of clonotypes identified increases with the fraction of CDR3 reads in the RNA-seq dataset (x-axis). **(f)** The T-cell clonal diversity, determined by the number of clonotypes divided by the normalized CDR3 read count, is similar in transcriptome (x-axis) and exome (y-axis) datasets. Two-dimensional density is represented (blue lines), as well as eleven tumors sharing clonotypes between DNA and RNA (red dots).

The majority (54%) of tumors with CDR3 reads in the exome displayed only one clonotype sequence and the number of clonotypes identified increased with the fraction of CDR3 reads identified (Fig. 4D). Indeed, in contrast to deep repertoire sequencing, our approach is not deep enough to saturate the T-cell clonal diversity and thus provides only a shallow view of the repertoire. Similarly, the number of clonotypes identified in the transcriptome data increased with the fraction of CDR3 reads (Fig. 4E). Importantly, the ratio of clonotypes to the normalized CDR3 read count could be used to approximate clonal diversity. This measurement had a large variance but was consistent between DNA and RNA (Fig. 4F, r=0.11, p=0.02). We observed a total of 839 and 7,130 different clonotypes across all tumors using the exome and transcriptome data, respectively, and eleven patients shared at least one clonotype between their tumor RNA and DNA. Oligo-clonality of the TCR repertoire could increase the chances of observing shared clonotypes between RNA and DNA of the same tumor, especially at shallow depth. However, none of these 11 tumors had noticeably low clonotype diversity (Fig. 4F), in agreement with the substantial under-sampling of the *TCRB* repertoire.

We next identified clonotypes shared between patients’ exomes (Supplementary Table 3). Two DNA clonotypes were shared between in 2 and 66 tumors, respectively. The most shared clonotype (66 tumors, referred to as c66) was also identified in the blood DNA of 36 of these patients and of an additional 40 patients. This suggests that the c66 clone was not tumorspecific, and may be directed against an antigen present relatively frequently in the population. Importantly, we did not find any significant association between the presence of the c66 clone and the patient HLA type (Supplementary Table 4), indicating that this TCR clone is likely reacting to a promiscuous antigen. Similarly, we did not identify a specific association of the c66 clone in patients with mutations in *PIK3CA*, *GATA3*, or *TP53*, the most common breast cancer driver genes (Supplementary Table 5).

## Discussion

The study of the tumor immune-environment is particularly challenging given the complexity of the immune response and of the variety of host-tumor interactions. Furthermore, there is a critical lack of immune-specific molecular and histological observations in large cohorts of human samples. While the immune response is highly patient-specific, large cohort studies can nevertheless inform on the global dynamics and diversity of the immune response. In this report, we used the breast cancer cohort from the TCGA, which is the most comprehensive molecularly annotated breast cancer cohort, to characterize further their immune-microenvironment.

Our study is complementary to the analysis of immune-gene expression signatures and presents a method to characterize T-cell infiltration directly from the bulk tumor DNA. We were specifically inspired by similar strategies using “junk” or unaligned reads from genome-wide tumor sequencing to identify non-canonical information such as mitochondrial DNA sequence^29^, telomere length^30^, microsatellites^31–33^, pathogens^34^ or B-cell repertoire^35^. In order to detect CDR3 reads in bulk exome or transcriptome sequencing, we used existing algorithms that were designed to analyze TCR targeted sequencing, either by multiplex PCR or RT-PCR.

Importantly, they all used a pre-defined set of reference V, D, and J gene combinations, together with local re-alignment and error correction. The tethering to the current reference TCR gene annotation may be limiting the sensitivity of the approach, including the exome approach we used, and de novo assembly of fully rearranged TCR genes may be preferred but has yet to be reliably implemented. We think, however, that our approach has a high specificity since 72% of the CDR3 reads detected in the exome were present in the deep TCR repertoire, which was generated using an independent PCR based method.

We show the clinical utility of the resulting iDNA score in Her2+ breast cancer. This result is consistent with previous reports^36^ and is likely due to the use of anti-Her2 therapy in most patients^37^, making the iDNA score a predictive rather than prognostic marker. Our analysis also suggests that it has higher value over global histological measurement of TILs in Her2+ patients. We did not find any prognostic value of TILs or iDNA score for TNBC or HR+/Her2-tumors, in contrast to some previous reports^4,37–39^. However, the cohorts used in these reports had much more complete clinical information and more refined study designs, including neoadjuvant treatment, tumor stage and recurrence, immunohistochemistry of different lymphocytes subtypes, or distinction between stromal and intra-tumoral TILs. None of these features are available or reliable in the TCGA cohort and therefore could not be used to replicate the results. The survival association obtained in Her2+ patients highlights the overall predictive value of infiltrating T-cells when antibody therapy is used. A similar approach could be used to interrogate TCGA data for the predictive value of immune profiles and T-cell infiltration for other monoclonal antibody therapies such as cetuximab in EGFR-mutated non-small cell lung cancer patients^40^.

Immuno-histochemistry and flow cytometry are traditionally used to characterize the tumor immune compartment and develop immune biomarkers^9^. However, these measurements typically lack the information about the clonal identity and diversity of the adaptive immune cells. Deep TCR repertoire sequencing has been used to study the relevance of clonal diversity in various cancers^18–22^. Low clonal diversity in the blood (divpenia) is associated with shorter survival in metastatic breast cancer patients diagnosed with lymphopenia^41^. The deep sequencing of several regions of a large clear cell renal carcinoma shows a high heterogeneity of the clonal cell distribution^20^. Furthermore, the TILs in colorectal adenocarcinomas are more oligoclonal (lower diversity) than the lymphocyte population characterized from non-adjacent normal mucosa from the same patient^18^, further suggesting that the diversity is dictated by the tumor biology and neo-antigen reactivity rather than by organ specific biology. Finally, the analysis of B-cell receptor (BCR) diversity through the TCGA RNA-Seq data showed evidence of oligo-clonality in basal-like and Her2-enriched breast cancer^147^. The tumor TCR repertoire identified in our study is shallow – 79% of tumors with fewer than 10 clonotypes from RNA or DNA – but broad – 7,954 clonotypes (RNA or DNA) identified in 944 patients. Importantly, we identify only 2 clonotypes in common between patients, suggesting the absence of a dominant public T-cell clonotype and underscoring the exquisite patient specific immune response in breast cancer. A previous deep repertoire study reported 29 out of 32,000 clonotypes shared between 2 of 15 colorectal cancer patients^18^, a proportion consistent with our observations in a shallow TCGA repertoire. Furthermore, it has been proposed that the clonal composition of lymphocytes differs between tumor, lymph nodes and in the circulation, the repertoire being locally shaped by the presence of tumor specific antigens^20^. Our data suggests that 7 T-cell clonotypes in 41 patients are also present in the circulation and therefore less likely to be directed at tumor specific antigens. This implies that the study of the tumor infiltrating T-cell repertoire requires the analysis of a synchronous matched blood or draining lymph node control to accurately identify tumor specific clonotypes, or determine their rate of tumor residency versus recirculation. Additional studies are needed to fully understand the regional variation of the repertoire and its consequences on cancer progression and response. Nevertheless, the value of TCR repertoire sequencing may be in monitoring the clonal evolution within a patient or tumor rather than in the identification of a broad-spectrum tumor specific antigen or its corresponding T-cell clone.

## Methods

### Deep T-cell repertoire sequencing

Matched frozen – formalin-fixed, paraffin-embedded (FFPE) tissue samples from three triple negative breast cancer (TNBC) tumors were obtained from Asterand Biosciences (Detroit, MI). These samples were selected for having mirrored specimens from FFPE blocks and fresh frozen tissue. A pathology inspection of hematoxylin and eosin (H&E) stained sections indicates that two tumors have histological evidence of TILs greater than 40%, while one sample was devoid of TILs as estimated and used as a negative control For each FFPE sample, we extracted DNA from ten 10 μM sections using the QIAamp DNA FFPE tissue kit (Qiagen, Venlo, Netherlands). The deep TCRB repertoire libraries were prepared using the ImmunoSeq kit (Adaptive Biotechnologies, Seattle, WA) following the manufacturer’s instructions. Briefly, 10 μg of DNA from each sample was split into 2 replicate (survey depth) to perform the 1^st^ PCR amplification (30 cycles). The entire product was then purified using the magnetic beads, eluted in 10 μL and subjected to the 2^nd^ PCR amplification (7 cycles). The resulting indexed libraries were pooled and sequenced in one run of a MiSeq using ImmunoSeq custom read1 and index primers and sequenced for 150bp (+6bp index read) using MiSeq reagents v3. The raw sequencing data was uploaded to Adaptive Biotechnologies’ server for analysis via the ImmunoSeq analyzer web application.

### Breast tumor exome sequencing

The sequencing libraries were prepared and captured using the SureSelect Human All Exon V4 kit (Agilent Technologies, Santa Clara, CA) following the manufacturer’s instructions. Briefly, 500 ng DNA was fragmented by Adaptive Focused Acoustics (E220 Focused Ultrasonicator, Covaris, Woburn, MA) to produce an average fragment size of ~175 bp and purified using the Agencourt AMPure XP beads (Beckman Coulter, Fullerton, CA). The quality of the fragmentation and purification was assessed with the Agilent 2100 Bioanalyzer. The fragment ends were repaired and adaptors were ligated to the fragments. The resulting DNA library was amplified by using the manufacturer’s recommended PCR conditions: 2’ at 98°C followed by 6 cycles of (98°C 30”; 65°C 30”; 72°C 1’) finished by 10’ at 72°C. 500 ng of each library was captured by solution hybridization to biotinylated RNA library baits for 48 hrs. at 65°C. Bound genomic DNA was purified with streptavidin coated magnetic Dynabeads (Invitrogen, Carlsbad, CA) and further amplified to add barcoding adapters using manufacturer’s recommended PCR conditions: 2’ at 98°C followed by 12 cycles of (98°C 30”; 57°C 30”; 72°C 1’) finished by 10’ at 72°C. The library was sequenced on one lane of HiSeq 2500 (PE 100pb reads). The resulting reads were mapped to hg19 using bwa-mem^42^ and duplicate reads were removed using Picard MarkDuplicates^43^.

### Identification of CDR3 reads

The reads mapped to the human TCRB region (hg19: chr7:142,000,817-142,510,993) and the unmapped reads were extracted from the BAM files, and converted to fastq using SAMtools^44^ and BEDtools^45^. The tools ClonotypeR^25^, MiTCR^27^, and IMSEQ^26^ were used to identify rearranged TCR regions from the resulting files. Human TCR references were provided by MiTCR and IMSEQ. We created our own reference for ClonotypeR as specified in the documentation. In short, we downloaded reference sequences for TCRA, B, and G from GenBank^46^, manually aligned the V and J segments on the conserved motifs using SeaView^47^, and separated out the segments from before or after the conserved motifs. We compared the performance of the three tools, matching on read ID. When the read ID was not available from the results (MiTCR), we retrieved it by aligning all reads (BWA) to a reference consisting of MiTCR output reads. For each tool, the number of CDR3 reads detected was normalized by the total number of reads sequenced for that sample.

### TCGA sample selection and data access

We retrieved the complete genomic data and clinical data from TCGA^48^. We selected patients with both: 1) primary tumor (no metastases) exome BAM files aligned to GRCh37-lite (for multiple versions, the latest was kept), and 2) known fraction of TILs. There were a total of 1078 breast cancer patients following these criteria. We also retrieved, for the same patients, when available: 3) blood exome BAM files aligned to GRCh37-lite, 4) RNA-seq tumor BAM files aligned to hg19, 5) known ER, PR, and Her2 status, 6) vital status/know days to last contact/days to death. The clinical data was retrieved from TCGA data portal^48^ for BRCA on 4/27/15. Survival analysis was done using the R “survival” package^49,50^. The sequencing data was accessed from the CGHub^51^. The normalized tumor gene expression values and the gene mutational status were retrieved from Broad Institute Firebrowse^52^. *TCRB* expression was missing from the FireBrowse datasets. In order to evaluate expression from *TCRB,* we retrieved reads mapped to the *TCRB* gene region from RNA-seq data for each patient from CGHub. We extracted the number of reads mapped to this region, and normalized by the total number of reads to calculate a *TCRB* expression value.

### Gene set enrichment analysis

We used the LM22 gene signature matrix^13^, which defines gene sets for 22 immune cell types, totaling 547 genes. We used Gene Set Variation Analysis (GSVA)^53^ to evaluate enrichment of these signatures in each sample using the Firebrowse expression data. This analysis was performed in R using the “gsva” package^53^. Clusters were defined by hierarchical clustering of the patients by their LM22 enrichment scores.

### HLA haplotype calling

Blood normal whole-exome sequencing data was downloaded from TCGA for the BRCA cohort. HLA class I types were identified through a consensus approach of three tools: optitype^54^, athlates^55^, and snp2hla^56^. Allele assignments were selected for cases when two or three tools agreed, and when the tools did not agree, alleles were assigned by optitype, as it has the highest reported accuracy^54^. HLA class II types were identified by merging the results of athlates and snp2hla, since each covered a different subset of the genes.

### Simulations of CDR3 read detection

The ratio of the length of the CDR3 VDJ region (50 bp) to the total length of a captured exome (50M bp) can be used to estimate the sensitivity to detect CDR3 reads in an exome sequencing experiment. The parameters used were 1) off target ratio of 20% (fraction of reads mapped outside of the exome) and 2) a CDR3 detection sensitivity of 50%. This latter parameter is derived from the fraction of *TCRB* exons captured by the Agilent SureSelect v4 (69/107 based on Gencode v19 annotation^57^, as well as imperfect mapping (fraction of reads spanning CDR3 detectable by IMSEQ depends on the length of the reads and the position of CDR3 reads across the VDJ junction).

## Acknowledgements

We very much appreciated the assistance of Dr. Kristen Jepsen and Mrs. Mahdieh Khosroheidari from the UCSD-IGM genomics center and of Dr Jack Bui for his review of the manuscript. The work was performed on the iDASH compute cloud which is supported by NIH grant U54HL108460 and UL1TR000100 to Dr. Ohno-Machado. OH is being supported by NCI grants 1R21CA177519 to Dr. Howell and Harismendy, 2P30CA023100 to Dr. Scott Lippmann and 1U01CA196406 to Dr. Laura Esserman. EL is supported by NLM grant T15LM011271 to San Diego Biomedical Informatics Education & Research (SABER). VGC was supported by an administrative supplement to NIH grant 2P30CA023100 and FONDEF D11I1029 from CONICYT-Chile. HC and RM are supported by NIH grant DP5OD017937. RM is supported by the NSF graduate fellowship award #2015205295.

## Author Contributions

The manuscript was written by EL and OH. Deep T-cell and tumor exome sequencing was done by BW, and analyzed by EL and OH. Identification of CDR3 reads in TCGA and data collection was done by EL and VGC, and analyzed by EL and OH. HLA haplotypes were called by RM and HC, and analyzed by MD.

---

**Supplementary Figure 1:**
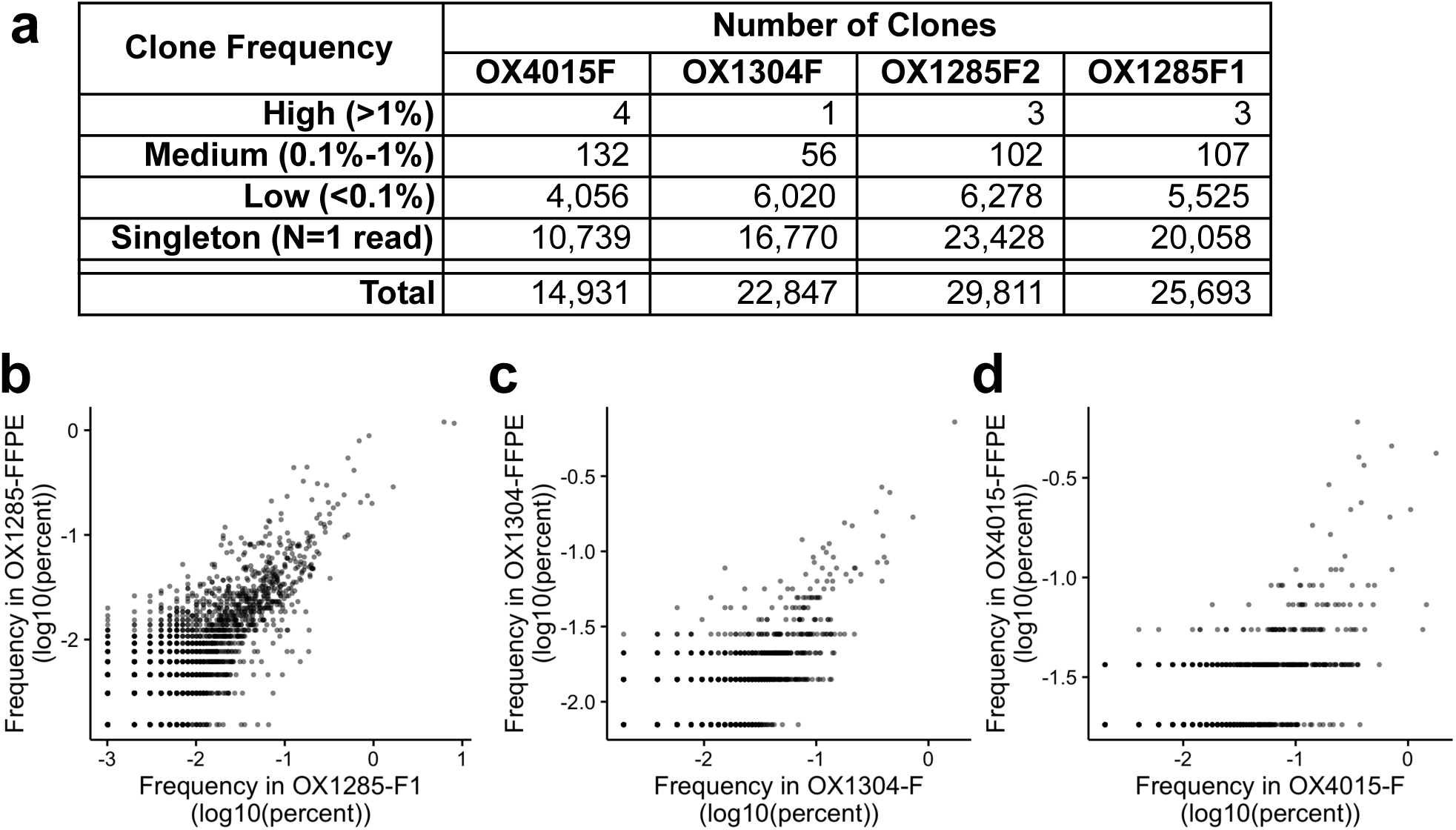
Deep Repertoire Sequencing by Adaptive ImmunoSeq. **(a)** Distribution of clone frequency in the four specimen studied. **(bd)** Scatter plot of clonotype abundance for mirrored frozen and FFPE tissue sections for samples OX1285 (b), OX1304 (c), 0×4015 (d).

**Supplementary Figure 2:**
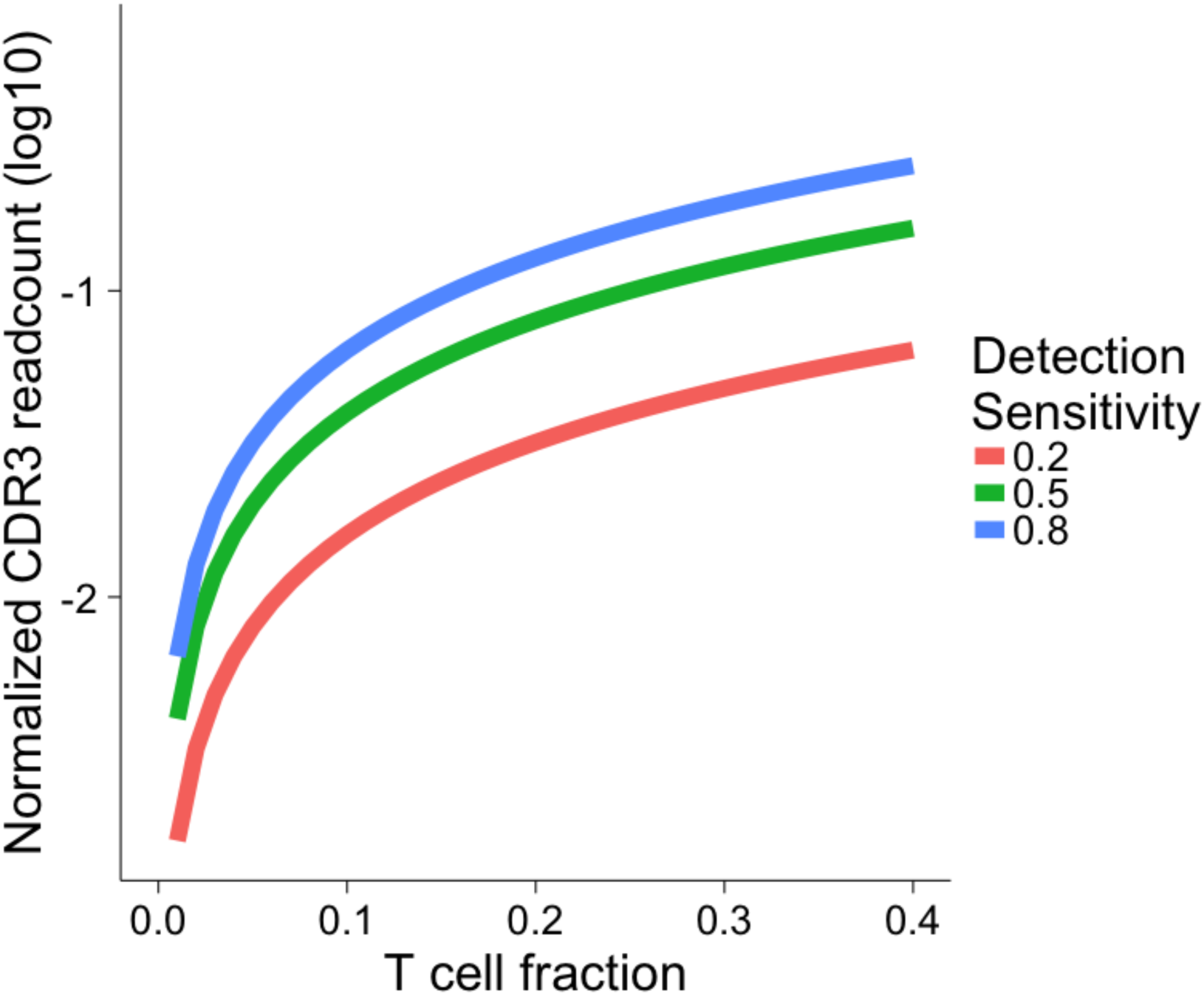
Read detection simulation for CDR3 reads in whole-exome data. Simulated fraction of CDR3 reads (y-axis) expected from a whole-exome sequencing experiment, as a function of T-cell fraction (x-axis) and CDR3 detection efficiency. See methods for details.

**Supplementary Figure 3:**
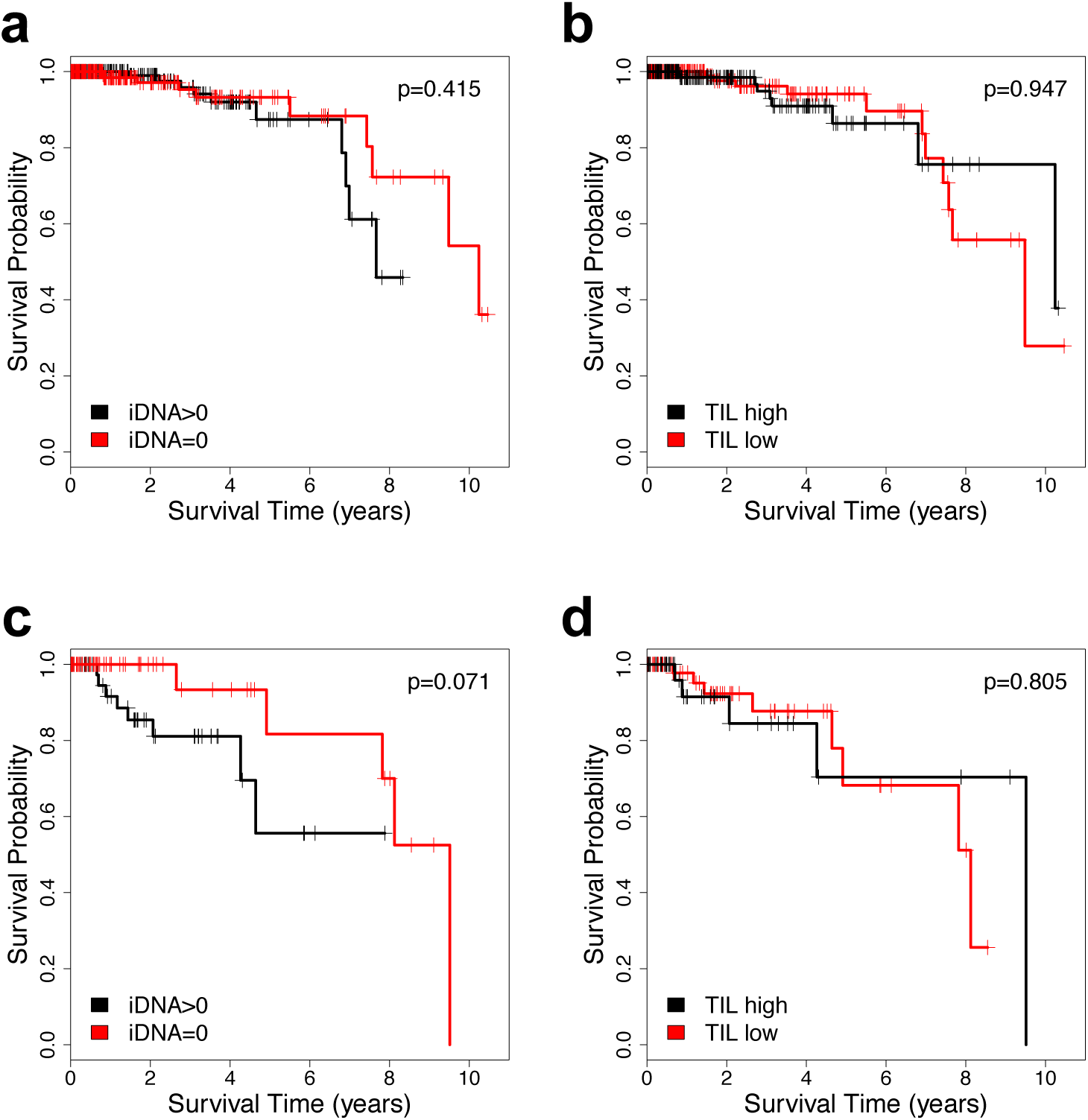
BRCA subtype iDNA and survival analysis. Kaplan-Meier survival analysis with significance of the hazard ratio of **(a**) HR+ patients as a function of iDNA score. Hazard-ratio is 0.676 [0.264-1.73]. **(b)** HR + patients as a function of TILs. Hazard-ratio is 1.03 [0.420-2.53]. **(c)** TNBC patients as a function of iDNA score. Hazard-ratio is 0.277 [0.069-1.12]. **(d)** TNBC patients as a function of TILs. Hazard-ratio is 0.859 [0.256-2.88].

